# Hub Façade: View Track Hubs in Integrated Genome Browser

**DOI:** 10.64898/2026.03.26.708838

**Authors:** Nowlan H. Freese, Karthik Raveendran, Jaya Sravani Sirigineedi, Udaya Chinta, Philip Badzuh, Omkar Marne, Chirag Shetty, Irvin Naylor, Saideepthi Jagarapu, Ann E. Loraine

## Abstract

**Summary:** Genome browsers are essential for understanding genomic data in detail but use incompatible data repository formats and access protocols, limiting their use. We present a new web application Hub Façade that translates the Track Hub format used by the UCSC Genome Browser into the Quickload format used by the Integrated Genome Browser (IGB), and vice versa. We used the Façade’s translation capability to write a single-page web application for IGB users to search and add UCSC-curated Hubs to IGB for exploration and visual analysis. We created a new IGB App (GenArk Genomes) that uses the Façade to automate importing genome assemblies from GenArk, a Hub-based repository with nearly 50,000 genomes hosted by the UCSC Genome Browser team. An example use case investigating alternative splicing of human gene *MEOX1* shows how using both browsers to view the same data via the Hub Façade promotes understanding and discovery.

**Availability and Implementation:** Hub Façade is free, open-source software deployed at translate.bioviz.org. Code is available from git repositories at [bitbucket.org|github.com]/lorainelab/hub-facade. The use case is available as Supplemental File 1.

## Introduction

All genome browsers display genomic data in the context of a reference genome assembly, but their implementations and capabilities vary. The University of Santa Cruz (UCSC) Genome Browser is a web site that offers searchable access to thousands of genome assemblies and curated data sets (Perez, et al., 2025). In contrast, the Integrated Genome Browser (IGB) is a compiled desktop application written in Java that users download, install, and run locally. The IGB design emphasizes visual analytics (Freese, et al., 2016; Gulledge, et al., 2014), interactivity (Nicol, et al., 2009), and extensibility via pluggable Apps (Shanbhag, et al., 2022). Locally installed IGB runs entirely on the user’s personal computer but can display data from on-line repositories alongside private data stored locally. Local endpoints within IGB can connect it to data provider web sites like BioViz *Connect* (Raveendran, et al., 2022) and eFP-Seq browser (Sullivan, et al., 2019). The user experience could not be more different between the two browsers, but they confront a universal problem: how to deliver data and reference genome assemblies to the software for visualization.

One way the UCSC Browser solves this problem is via a data repository format called Track Hubs (Raney, et al., 2014). Track Hubs are configuration files stored on-line in locations visible to the UCSC server that reference dataset files (tracks) via URLs. To open and view a Track Hub in the UCSC Browser, users enter its URL on a page at the UCSC site. The site retrieves format-specified configuration files from the provided URL and shows them alongside already-available native tracks. Users can create and share Track Hubs with the public via a Public Track Hub page on the UCSC genome browser site. The UCSC team also uses the format to distribute thousands of reference assemblies as part of the GenArk project (Clawson, et al., 2023).

Preceding Track Hubs, the IGB team developed IGB Quickload, based on a similar idea (Nicol, et al., 2009). Quickloads consist of REST endpoints that deliver information about where data files are located and how they should look once opened. To add a Quickload to IGB, users open a “Data Sources” configuration window and enter the URL or file path leading to the Quickload’s root directory. IGB retrieves configuration files from the Quickload site and uses them to build interfaces for users to select, display and configure assemblies and data tracks. Today’s IGB team uses internet-accessible Quickload sites to provide diverse reference data sets and assemblies.

Quickload and Track Hub configuration files use simple, text-based formats, requiring minimal or no programming skill to create. However, neither browser can read the other’s configuration files. If someone wants to use both browsers, they must set up and maintain two repositories, one for each browser. Few groups do this, even though using multiple browsers with complementary capabilities and strengths can yield important discoveries. The technical similarities between the formats suggest a solution. Using software, we should be able to translate configuration files on demand, giving IGB access to data previously visible in the UCSC Browser only, and vice versa.

## Results

We created Hub Façade, a web application that dynamically converts Track Hub configuration files to their Quickload counterparts, and vice versa, enabling visualization of the same data in Integrated Genome Browser and the UCSC Genome Browser at the same time. We used the Façade’s translation capability to implement two new IGB data access resources. One is a single-page web application listing public Track Hubs with controls for opening the Hubs in IGB. The second is a new IGB App that automates translating UCSC GenArk Track Hubs into IGB Quickloads, making thousands of assemblies readily available in IGB for the first time. To illustrate the value of this work, we present an example use case investigating alternative splicing frequency of human gene *MEOX1*.

### Track Hub Façade web application

We deployed the Hub Façade to translate.bioviz.org, using ansible playbooks available from the Façade git repository. The deployed application displays a web page front end where users can manually enter Track Hub URLs to obtain an IGB Quickload equivalent. **Figure 1a** shows an example where a user has entered a track hub URL in the upper input field and clicked the “Convert” button. JavaScript loaded with the page has produced an IGB Quickload URL that references translate.bioviz.org and specifies the Track Hub source using parameter hubUrl. Clicking the button labeled “Add to IGB” imports the new Quickload into IGB using a localhost endpoint implemented within IGB. Clicking the vertical double arrow switches to translating from Quickload to Hub format.

**Figure 1.**
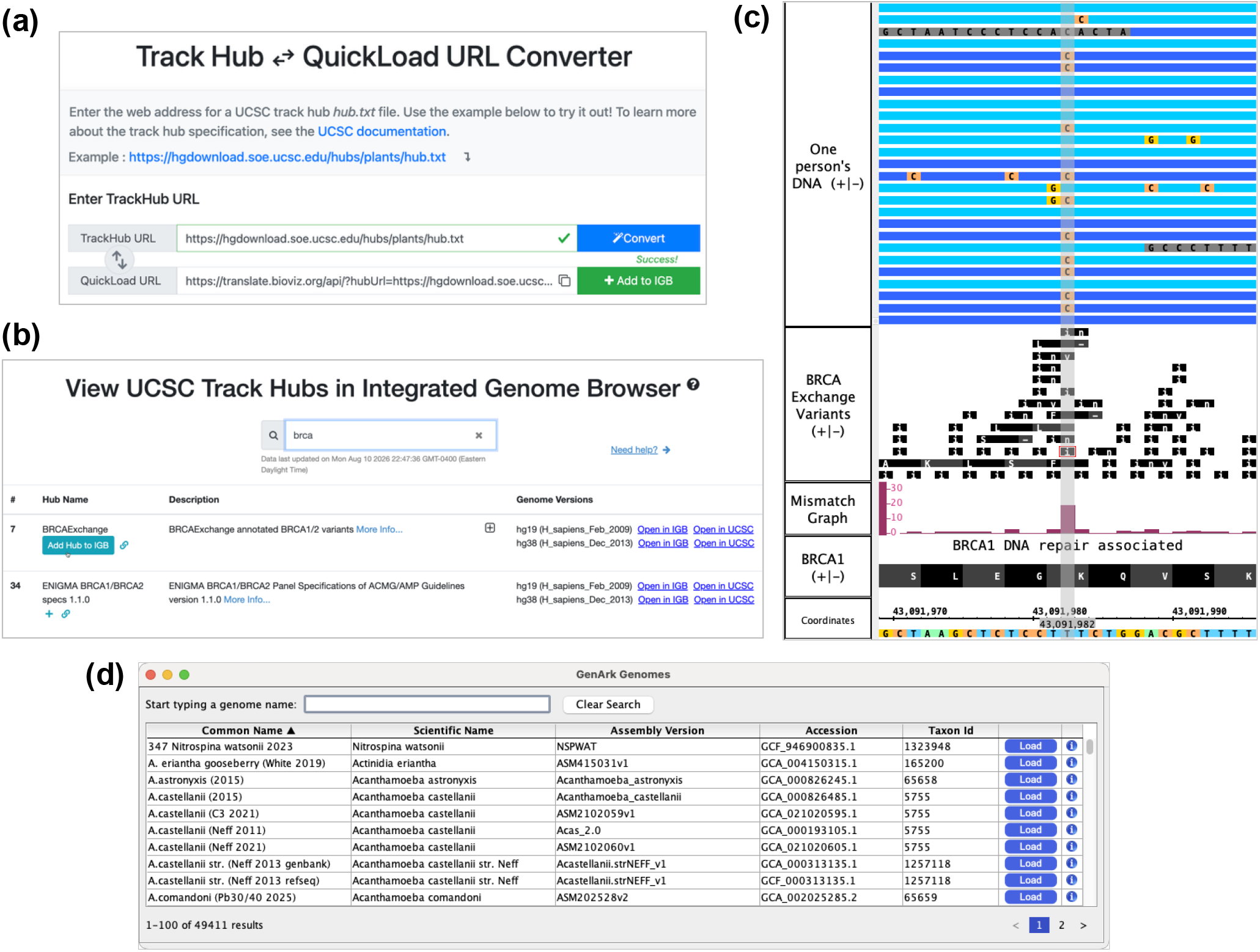
**(a)** Hub Façade web interface showing a UCSC track hub URL converted to an IGB Quickload URL. Clicking “Add to IGB” button adds the Quickload URL to a locally running IGB instance. **(b**) Bioviz.org Public hubs page after searching for tracks that match query string brca. (**c**) Image from IGB where a user has loaded genome sequencing data alongside a variants track imported from the BRCA Exchange Track Hub. (**d**) Snapshot of GenArk Genomes app tabbed panel running in IGB showing UCSC GenArk assemblies available to open in IGB

### Public Track Hubs page

The UCSC Genome Browser web site maintains an API (application programming interface) that provides programmatic access to a listing of Track Hubs that research labs and organizations have created and made available to the public. Using the API, we wrote a single-page web application that displays a searchable table of these public Hubs (**Figure 1b**). Within the table, rows contain user interface components for importing a given Hub into IGB or the UCSC Browser. **Figure 1c** shows an image from IGB where a hypothetical user has added a track from the first search result, a Track Hub from the BRCA Exchange (Cline, et al., 2018). The user has also imported alignments of their own genomic DNA into a track and created a mismatch graph summarizing differences between their sequence and the reference, shown in the track labeled “Mismatch Graph”. Using the mismatch graph and the alignments track, they can identify variants in their own DNA, match them to variants in the BRCA Exchange track, and then look up their possible clinical significance using the BRCA Exchange web site.

### GenArk Genomes IGB App

The UCSC Genome Archive (GenArk) (Clawson, et al., 2023) makes thousands of genome assemblies accessible via the Track Hub format. To make it easy for IGB users to find and load these assemblies within IGB, we created GenArk Genomes, a new IGB App, and released it on the IGB App Store web site (Shanbhag, et al., 2022). Once installed into IGB, the GenArk App adds a new tabbed panel interface to IGB (**Figure 1d**). Users can search for assemblies by species common name, scientific name, assembly version, accession, or taxon id. When a user selects an assembly and clicks the load button, the App contacts the Track Hub Façade backend API to obtain a Quickload URL and adds it to IGB using a localhost endpoint implemented within IGB. The selected genome then opens in IGB. Associated data tracks, such as gene annotations, appear in the Available Data section of the IGB Data Access tab. From there users can load data and utilize IGB’s visual analytics tools as if the genome assembly were provided by a native IGB Quickload.

### Example use case

We created a Track Hub with RNA-Seq alignments from twenty human tissues to investigate alternative splicing of a human gene encoding a DNA-binding, regulatory protein (MEOX1), a follow-up study from previous work (Loraine, et al., 2003). Using the Façade to view the same data in both IGB and the UCSC Browser, we found that an exon-skipped splice variant lacking a conserved DNA-binding domain was always expressed at very low levels compared to the non-skipped form. The skipped exon’s donor site coincided with base positions flagged by AbSplice, software that predicts tissue-specific, aberrant splicing (Supplemental File 1.)

## Methods

Hub Façade is implemented using the Django web application framework. The site uses a façade pattern (Gamma, et al., 1994), presenting an API that mimics expected endpoints for the target repository format. For example, Quickload requires an “annots.xml” endpoint that lists available data sets, URLs for loading the data, and visual configurations. Likewise, Track Hub requires a “trackDb.txt” endpoint with similar information. A converted Hub to Quickload URL or vice versa has parameters “hubURL” or “quickloadURL” indicating desired direction of translation and the source repository.

For the example use case, we aligned RNA-Seq sequence reads from Sequence Read Archive Study SRP056969 (BioProject PRJNA280600) to reference human genome assembly hg38 and deployed the resulting data files on-line in various locations. We then manually created Track Hub configuration files that reference the data file URLs. We deployed the files on-line at https://bitbucket.org/nfreese/trackhub-human-hub. Instructions for loading the Hub into the UCSC Browser or IGB are available there in a README. To view the data, we used Integrated Genome Browser version 10.2.0 and UCSC Genome Browser version(s) available in summer of 2026.

## Discussion

The Track Hub Façade dynamically translates Track Hub configuration files into endpoints provided by an IGB Quickload site. Using the Façade’s back-end API, we created two new software applications to facilitate loading data from Track Hub repositories into IGB. The first was a single-page JavaScript application (https://bioviz.org/public-trackhubs.html) that parses Hubs listed at https://genome.ucsc.edu/cgi-bin/hgHubConnect and creates an interface that automates adding the new Hubs to IGB. The second was a new IGB plugin App “GenArk Genomes” that connects IGB to thousands of reference genome assemblies hosted as Track Hubs on UCSC Genome Browser web resources. Thanks to this new work, IGB users can now open and explore existing Track Hubs contributed by many different groups. The relative privacy IGB offers, along with its greater extensibility compared to other genome browsers, expands access to audiences with diverse needs and interests.

After implementing the Hub to Quickload translation functionality, we added a newer capability to convert Quickload repositories to Track Hubs, so that our existing IGB Quickload sites could be shown in the UCSC Browser. In the process, we faced two main challenges. First, IGB accepts some data file formats not recognized by the UCSC Browser, notably, tabix-indexed files (Li, 2011). This means that many files offered in a Quickload site cannot be shown in the UCSC Browser. Second, the Track Hub specification sometimes requires specifying file type, but the Quickload specification does not. Thus, the Quickload specification lacks information the UCSC Browser requires for proper functionality. Future work may address these limitations.

## Supporting information

Supplemental File 1 - MEOX1 alternative splicing

Supplemental File 2 - MEOX1 ProtAnnot results

## Author contributions

NHF (Conceptualization, Supervision, Writing – original draft), KR (Software, Writing – original draft), JSS (Software), UC (Software), PB (Software, Writing – original draft), OM (Software), CS (Software), IN (Software), SJ (Software), AEL (Conceptualization, Supervision, Writing – review & editing)

## Acknowledgements

We thank Maximilian Haeussler and Luis Nassar for technical discussions regarding the track hub format.

## Supplementary Data

ProtAnnot data file with InterProScan search results is available on-line as Supplementary Data File 1.

## Data Availability

Links to data from the RNA-Seq alternative splicing use case can be found in hg38/trackDb.txt from https://bitbucket.org/nfreese/trackhub-human-hub.

