## Supplemental File 1 - MEOX1 alternative splicing for "Hub Façade: View Track Hubs in Integrated Genome Browser"

### Introduction

This Supplemental File describes a visual analysis use case demonstrating the Hub Façade Quickload Translator Web application, together with Integrated Genome Browser (IGB) (Freese, et al., 2016) and the UCSC Genome browser (Perez, et al., 2025). In this example, we investigated alternative splicing of the human gene mesenchyme homeobox 1 (*MEOX1*).

According to the National Center for Biotechnology Information (NCBI) Gene database (Sayers, et al., 2025), *MEOX1* is “a member of a subfamily of non-clustered, diverged, antennapedia-like homeobox-containing genes.” The Gene database record reports multiple distinct gene models, one of which skips a coding region exon. Loss of this second exon causes a frameshift in the third, downstream exon, deleting the protein’s DNA-binding region, a homeobox domain (Loraine, et al., 2003).

Loss of DNA-binding coding sequence in the exon-skipped variant suggests that this alternatively spliced transcript encodes an inactive, non-functional protein. However, if the skipped form were to be expressed at a high level, it could interfere with or suppress non-exon-skipped *MEOX1* activity by sequestering *MEOX1* protein binding partners, a dominant negative relationship. Thus, to understand the function of the gene in regulating target gene expression, it is essential to determine typical expression levels of exon-skipped versus non-exon-skipped variants. If the exon-skipped form is lowly expressed, it is likely to have little or no effect on *MEOX1* function. To answer this first question, we will use the Integrated Genome Browser, taking advantage of its visual analytics capabilities. Depending on the results, the next step will be to investigate how *MEOX1* splicing may be regulated. For this latter question, we will use the UCSC Genome Browser, taking advantage of its vast collection of data sets from experimental and computational studies.

### Results

To investigate relative expression levels of splice variants, we used an RNA-Seq data set from the Sequence Read Archive, available as study SRP056969, BioProject PRJNA280600. The data set contains sequences from twenty healthy human tissue types and was released as part of a study investigating recursive splicing (Duff, et al., 2015). We downloaded and aligned the sequences against an hg38 reference human genome assembly, uploaded the output files to a server for on-line hosting, and created Track Hub configuration files that organize the data files into a Track Hub. We deployed the Track Hub configuration files using free hosting within a Bitbucket git repository (https://bitbucket.org/nfreese/trackhub-human-hub). We then used the Hub Façade's Hub to Quickload translation feature to open the hub in Integrated Genome Browser and answer: Is the exon-skipped form highly or lowly expressed compared to the exon-included form?

Figure 1a shows the Track Hub loaded into IGB via the Track Hub Quickload Translator. The track labeled “Heart_reads” (from the Track Hub) shows RNA-Seq read alignments from heart, the sample with the most alignments in the *MEOX1* gene region. The vertical dimension within the track shows fifteen distinct rows of individual alignments, with a top row showing a squished, summary view of the remaining alignments, too numerous to be shown individually. The track below it labeled “RefSeq Curated (+/-)” shows three reference *MEOX1* gene models, each appearing as a series of linked blocks, where blocks represent exons and links represent introns removed during splicing. The “<” symbols superimposed on the intron connector links signal that transcription proceeds from right to left, using the plus (visible) strand of the reference assembly sequence as a template.

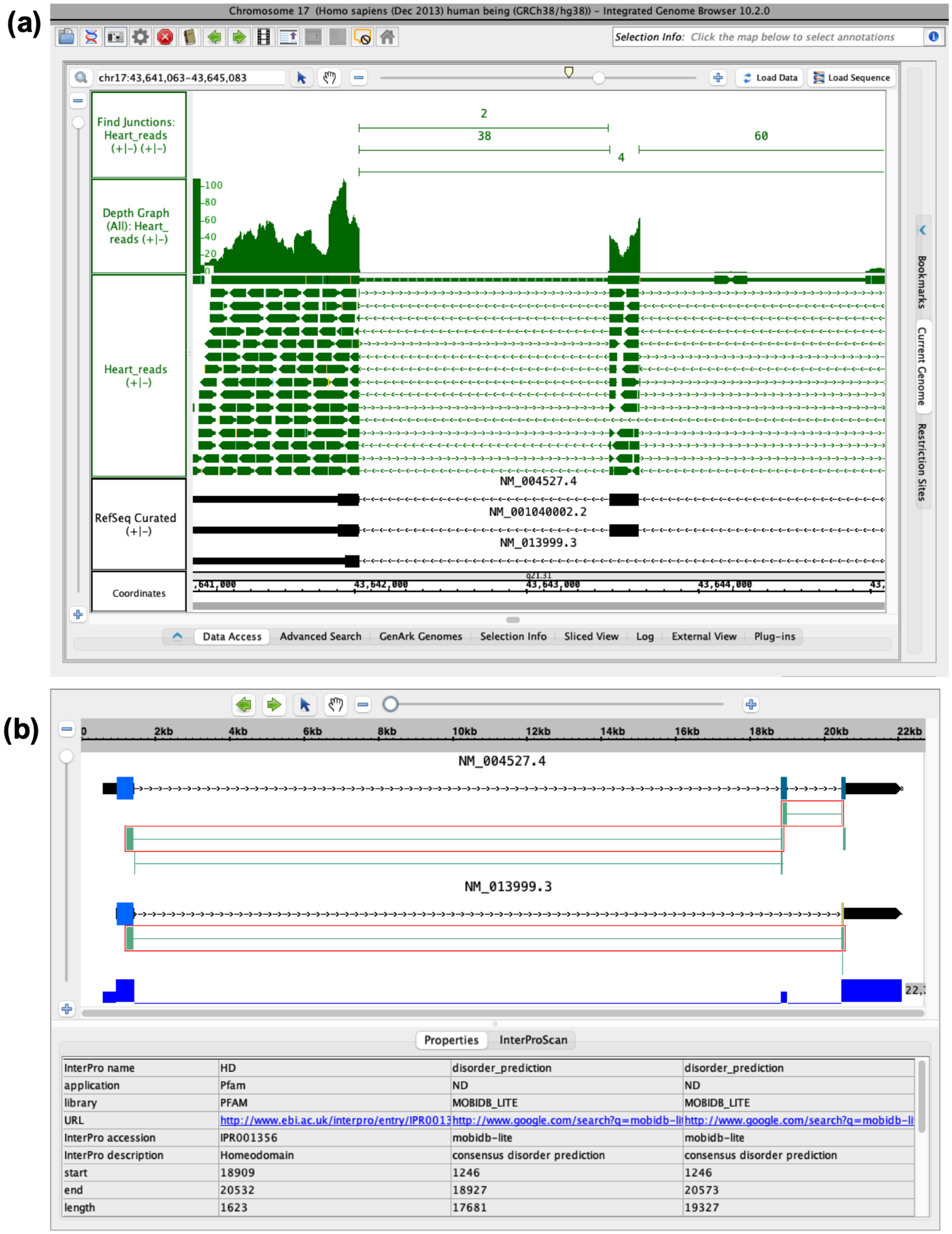


**Figure 1. Example Track Hub loaded in the Integrated Genome Browser (IGB) showing RNA-Seq alignments from heart tissue to the gene *MEOX1*. (a)** IGB interface displaying tracks from top down: Find Junctions track generated by IGB with counts of reads split by introns, Depth Graph generated by IGB displaying read counts, heart RNA-Seq reads aligned to genome assembly hg38, and *MEOX1* gene models. **(b)** ProtAnnot showing *MEOX1* gene models and selected InterProScan predicted protein motifs (green blocks outlined in red)

To assess RNA-Seq evidence for the exon-skipping splicing pattern, we used two IGB track operators, functions that make new tracks from existing tracks by applying visual analytics algorithms. These were: “Find Junctions” and “Depth Graph” (Gulledge, et al., 2014). The Find Junctions operator compares spliced RNA-Seq alignments within a selected track to each other to create non-redundant junction features capturing every splicing pattern evident in the data. To show the results, it creates a new track of exon-intron-exon features summarizing splicing evidence from the source track. Each exon-intron-feature in a Find Junctions track appears with a numeric label, a score, indicating the number of alignments in the source track with gaps matching the feature’s intron boundaries. The Depth Graph track operator summarizes the pattern and number of annotation overlapping a region. It makes a new graph track in IGB from a user-selected source track, in this case the RNA-Seq alignments track. It counts the alignment blocks overlapping each base position along the horizontal sequence axis, the “Coordinates” track number line and displays the result in the graph. (Another type of depth graph available in IGB counts only the first aligned base in per alignment.)

The track labeled “Depth Graph (All): Heart_reads (+/-)” in Figure 1a contains the Depth Graph operator output, with a y-axis on the left side of the track showing the number of alignments per base. Typical for RNA-Seq coverage graphs, the number of overlapping alignments varies across the gene body, as shown by the uneven peaks and valleys formed by the graph’s top edge. In this view, coverage coincides with gene model exons, but we also see two small, low coverage regions of RNA-Seq alignments in the 5’ (right most) intron. Visualization of the heart sample compared to the other nineteen tissues showed that the heart RNA-Seq data contained more *MEOX1* alignments than the others (not shown).

The Find Junctions track quantifies exon-exon junctions exposed by gapped sequence read alignments in the “Heart_reads” track, its source track. Comparing the RefSeq Curated track to the Find Junctions track shows that only four sequence read alignments from this dataset support the exon skipping event, compared to 38 and 60 alignments supporting exon inclusion, indicating that exon skipping was a rare splicing pattern in this sample. The Find Junctions track also exposed another rare splicing event where the right-side boundary of the downstream intron was eleven bases downstream of the more frequently used boundary. However, none of the annotated gene models matched this rare splicing pattern. We also examined *MEOX1* splicing in all other tissue samples (not shown). Only two other samples contained exon-skipped variants: prostate and lung. Of the three samples with exon-skipping, the SRP056969 prostate sample had the highest percentage of exon-skipped variant relative to the exon-included form (not shown). These results indicate that the exon-skipped form can occur in specific healthy human tissues but is expressed at lower levels than the fully functional, exon-included isoform.

Previously, we used an early, stand-alone version of the IGB App ProtAnnot to investigate functional consequences of exon-skipping in *MEOX1* transcripts (Loraine, et al., 2003). We found that the exon-skipped *MEOX1* splice variant deletes a part of a conserved homeodomain, suggesting it is non-functional (Loraine, et al., 2003). To check this prior result, we used a newer, updated version of ProtAnnot (Mall, et al., 2016), re-factored to function as an IGB plug-in, called an IGB App. This plug-in is available from the IGB App Store Web site (Shanbhag, et al., 2022) and from the App Manager menu option within IGB itself. After installing the App in IGB, users can select gene models and choose an option “Start ProtAnnot” from the IGB Tools menu or by right-clicking a selected gene model. From there, IGB opens a new ProtAnnot window showing the selected models, using exon fill color to indicate translation frame. By comparing exon colors vertically across stacked gene models, users can quickly notice frame-altering differences between splice variants. The app version of

ProtAnnot also enables users to search gene model conceptual translations against the InterPro compendium (Blum, et al., 2025) of conserved amino acid motifs and domains. Search results are then displayed graphically as green boxes next to the gene models, showing which exons or parts of exons encode conserved motifs.

Figure 1b shows an image created using this up-to-date version of ProtAnnot. In this view, ProtAnnot shows that exon-included gene model NM_004527.4 contains a homeodomain which is missing from the exon skipped form NM_013999.3. The difference in color for the final coding exon signals how exon skipping causes a frameshift in that exon. This visualization confirms the previous finding that the exon-skipped form lacks a homeodomain and likely cannot bind DNA, whereas the exon-included form probably can because it encodes an intact homeodomain region. We also note a new result that the amino terminal region of the protein, upstream of the homeodomain, contains a long intrinsically disordered region, detected by MobiDB-Lite (Mehdiabadi, et al., 2025). Long intrinsically disordered regions lack a single, stable folding structure and are thought to change conformations dynamically, depending on a protein's intra- or inter-molecular interactions (Aspromonte, et al., 2024). For clarity, we show only the results from searching MobiDB-Lite and Pfam, as the other database results were redundant with what is shown. A ProtAnnot file with results from searching all constituent databases using gene models NM_004527, NM_013999, and NM_001040002.2 is available as Supplemental File 2.

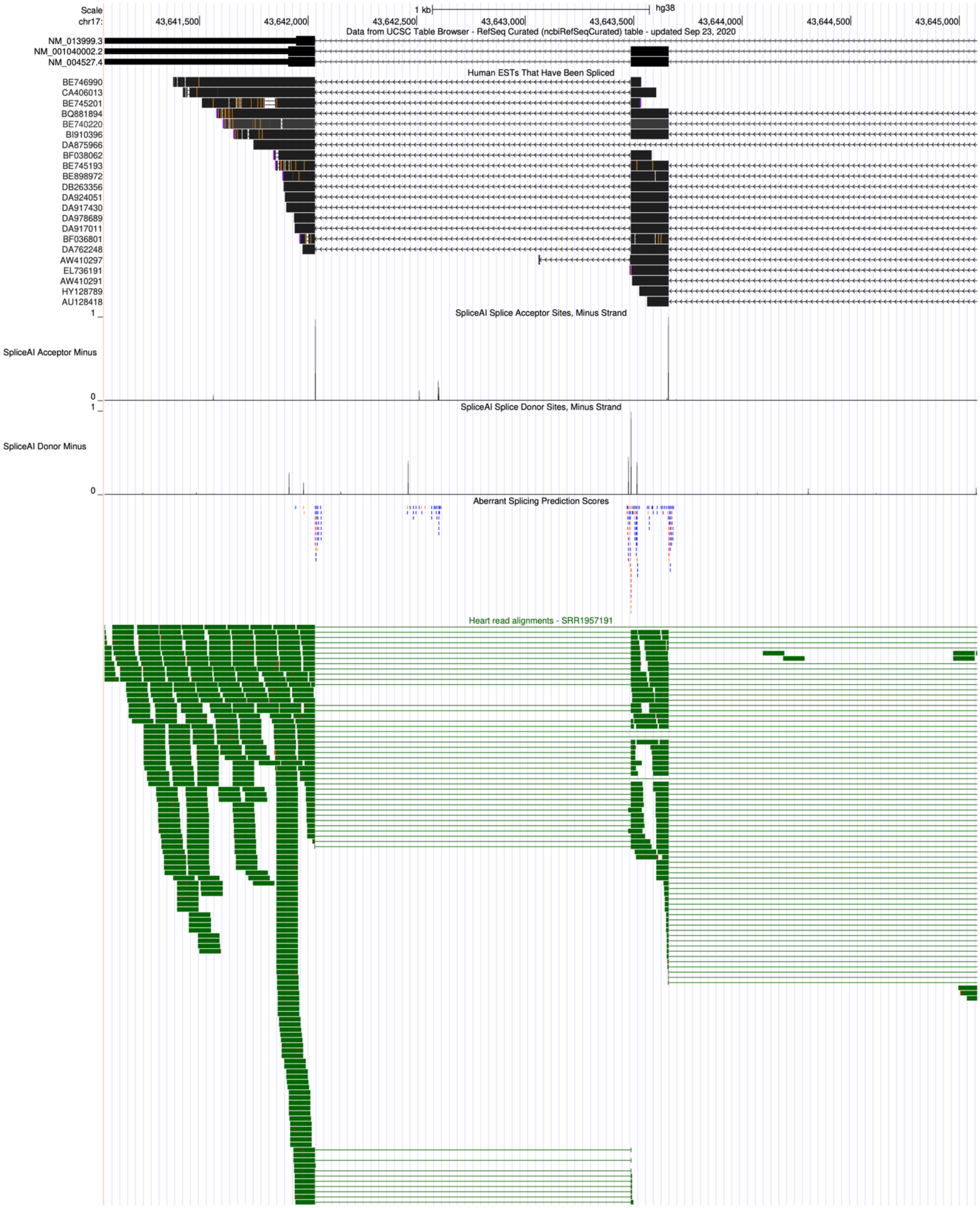


**Figure 2. UCSC Genome Browser showing human gene *MEOX1* in assembly hg38.** Tracks shown are *MEOX1* gene models (top), spliced expressed sequence tags (ESTs), three splicing related tracks, and RNA-Seq alignments from heart imported from the Track Hub created for this study.

The possible tissue-specificity of exon-skipping alternative splicing prompted us to investigate regulation of *MEOX1* splicing using the UCSC Genome Browser. The Browser offers access to thousands of tracks, including some relevant to splicing regulation. However, to use these tracks, we needed to be able to see the same RNA-Seq alignments imported into the browser for comparison. To do this, we added the Track Hub to the UCSC Browser, following instructions provided here: https://bitbucket.org/nfreese/trackhub-human-hub/src/main/. Once added, the Track Hub appeared as a new section on the browser page, allowing us to load and configure our data into the browser, which we did.

Next, we configured the browser settings to show five native UCSC data tracks relevant to splicing prevalence and regulation. These were: “NCBI RefSeq Genes,” “Human ESTs That Have Been Spliced,” “SpliceAI Acceptor Minus,” “SpliceAI Donor Minus,” and “Aberrant Splicing Prediction Scores.” These and other native tracks can be added to the genome browser’s current view by scrolling down the browser display page and selecting options from the sections below. The NCBI RefSeq genes track was from the section named “Genes and Gene Predictions” and the Spliced ESTs track was from the section labeled “RNA and Transcriptome. The other three tracks were from what the UCSC browser calls a “container track” labeled “Splicing Impact” in the section titled “Phenotypes, Variants, and Literature.”

Figure 2 shows an image from the UCSC Browser positioned to show the middle and terminal exon regions for the *MEOX1* gene after loading and configuring the above tracks. Three reference gene models appear at the top of the image, labeled with their RefSeq accessions. As in IGB, the UCSC Browser shows the direction of transcription for minus strand genes like *MEOX1* as proceeding from right to left. The RNA-Seq alignments from the Track Hub heart sample are shown in a tall track in the lower part of the image. In this view, the aligned sequences from the Track Hub are shown with the track in fully expanded mode, where every aligned sequence is visible in enough detail to observe individual exon boundaries. This differs from the IGB image, which we configured to show fewer distinct alignments so that we could also show the track operator outputs in the figure. In this expanded view, because each alignment is visible, we can count four alignments that support the exon-skipped form, matching IGB's Find Junctions results. We also see two alignments that appear to use a 3' splice (donor) site slightly to the left of the sometimes-skipped exon's usual boundary. These match IGB's Find Junctions result that showed a novel junction with a score of two.

The Spliced ESTs track is labeled "Human ESTs That Have Been Spliced" and is directly below the gene models track. (ESTs are "expressed sequence tags" that come from single-pass Sanger sequencing of cDNAs synthesized at random from RNA samples. These were used for many years to identify genes and gene structures in DNA.) According to the UCSC browser documentation, the track contains alignments for human-origin ESTs submitted to GenBank (Benson, et al., 2013) that contained evidence of at least one "canonical" intron, where "canonical" means: intron 5' and 3' boundaries start and end with bases GT and AG, and introns are at least 30 bases in size. Of the spliced ESTs that overlapped *MEOX1*, only one (DA875966) supported the exon skipped variant. Clicking the DA875966 EST in the browser display opened a popup with more information, revealing it was isolated and sequenced from prostate tissue as part of a large-scale EST (RNA) sequencing project (Kimura, et al., 2006). The other spliced ESTs supported the exon-included splicing pattern and came from diverse tissues.

The SpliceAI Acceptor Minus and SpliceAI Donor Minus tracks, below the ESTs track, came from running SpliceAI, an open-source deep learning algorithm that predicts splice site usage probability (Jaganathan, et al., 2019). Using the UCSC browser, we viewed minus strand, SpliceAI-predicted donor and acceptor sites, selected from the data set named “SpliceAI WildType.” (Donor sites mark the 3’ ends of exons and acceptor sites mark the 5’ ends of exons.) We observed that the SpliceAI tracks contained donor and acceptor sites corresponding to gap boundaries in the RNA-Seq, EST and gene tracks, along with several that did not.

SpliceAI predicted two additional donor sites in addition to the annotated donor site on the 3’ end (left side) of the sometimes-skipped exon. Zooming in for a more detailed view in IGB and the UCSC Browser, we found that one of these SpliceAI-predicted donor sites was supported by the two anomalous alignments mentioned previously, the same two alignments IGB’s FindJunctions track summarized as a single junction feature with score of 2. Using these alignments as a guide in IGB, we found that using this predicted, novel donor would add eleven bases to the exon boundary if used during splicing (Figure 3). Because eleven is not evenly divisible by three, it is likely to interrupt the reading frame, producing yet another, as-yet undocumented potentially defective *MEOX1* protein.

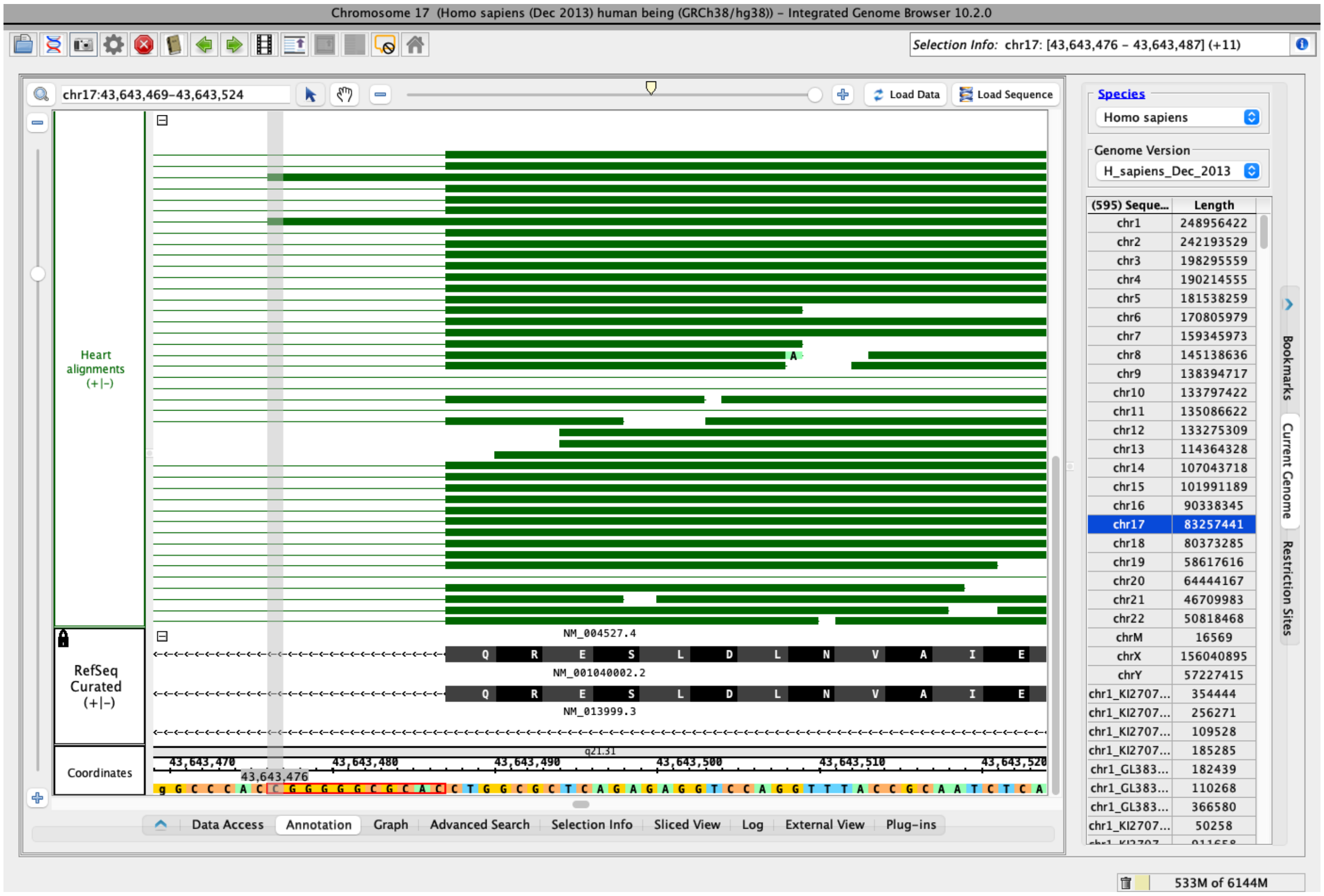


**Figure 3. Integrated Genome Browser image showing detailed view of the sometimes-skipped exon.** The user has click-dragged to select the genomic sequence bases extending beyond the exon's left boundary, and the upper right text box (labeled "Selection Info") reports the number of bases selected.

Next, we examined the AbSplice Scores track in the UCSC Browser. The AbSplice track shows single-nucleotide variants (SNVs) scored by their possible contribution to tissue-specific abnormal splicing (Wagner, et al., 2023). AbSplice variants appear as color-coded marks, where warmer (red) colors represent higher scores. Looking again at the second, sometimes-skipped exon, we observed a clustering of orange and red AbSplice scores around its donor and acceptor sites indicating a medium to high likelihood of aberrant splicing. Zooming in to select individual sites, we observed that the highest scoring sites were for tissues that included heart atrial appendage and heart left ventricle. In the SRP056969 data set, the heart sample had the most *MEOX1* alignments and also the most exon-skipped alignments. Thus, the AbSplice predictive result lined up with the experimental result.

### Discussion

We used a Track Hub data source and a website equipped with a browser-to-browser data source translator to view the same data in two genomes browsers with very different characteristics. As a test of the new browser-to-browser translator software and its ability to support new directions in research, we investigated alternative splicing of the gene mesenchyme homeobox 1 (*MEOX1*), with human genome assembly hg38 providing a common reference for alignment and comparison across data sets.

Using the IGB App ProtAnnot (Mall, et al., 2016), we re-confirmed an older result that exon-skipping deletes a DNA-binding homeobox domain encoded by the second and third *MEOX1* exons (Loraine, et al., 2003). We also observed something new: the amino terminal region, which was not affected by alternative splicing, contained an intrinsically disordered region, a protein sequence feature associated with flexible structures that can change depending on the cellular milieu (Aspromonte, et al., 2024). Such regions are also associated with protein-protein interactions (Aspromonte, et al., 2024), suggesting that the exon-skipped form may be able to interact with MEOX1 protein interactors despite lacking a DNA binding domain. This supports the hypothesis that the exon-skipped form, if translated into protein, could antagonize MEOX1 protein function.

Because our first investigation of *MEOX1* many years ago lacked access to RNA expression data, we could not determine the relative abundance of the exon-skipped versus non-skipped splicing variants. We also knew very little (at the time) about how DNA sequences could affect splicing rates and outcomes. In this new study, we used the Hub Façade's browser-to-browser translation function to import RNA-Seq alignments from a Track Hub into IGB, where we used IGB's analytical capabilities to investigate splicing frequency in SRP056969, an RNA-Seq data set from 20 different human tissues. Then, we opened and inspected the same data set in the UCSC Genome Browser, using its splicing-related tracks to investigate possible splicing regulation.

Our results included several important findings. First, we found that the exon-skipped form was lowly expressed relative to the exon-included form, the form that preserves the *MEOX1* DNA-binding domain. Second, we found that the exon-skipped form occurred in three data sets, and the relative abundance of the exon-skipped form versus the non-skipped form was highest in the SRP056969 prostate sample. We observed that the only EST in the UCSC Spliced EST track that supported the exon-skipped form was also from prostate, an intriguing coincidence. Third, we observed evidence for another splice form that adds 11 bases to the sometimes-skipped exon's carboxy-terminal end, introducing a frameshift and likely changing the C-terminal conceptual translation. This alternative splice variant coincided with SpliceAI and AbSplice annotations.

This latter finding was intriguing, but to understand its significance, we needed to know if the RNA-Seq data set we used (SRP056969) was included in SpliceAI or AbSplice training data sets. If the same datasets were used during training, any coincidences between what we observed in the SRP056969 data would mostly just show that the algorithms used to create SpliceAI and AbSplice models are good at detecting patterns in data. If not, the correspondence of results suggests a biologically meaningful trend or pattern, something reproducible across experiments and data sets. According to the documentation we were able to find in the browser and publications, training data included RNA-Seq alignments from the

Genotype-Tissue Expression (GTEx) collection, which, so far as we know, did not include SRP056969. If that is correct, then the AbSplice and SpliceAI algorithmic results suggest that MEOX1 splicing patterns we observed in SRP056969 are reproducible. As such, we can expect them to reappear in other data sets from new experiments.

The Hub Façade functioned as we had hoped, giving us a relatively friction-free ability to view the same data in our default browser (IGB), software we developed and know very well, and another, less familiar browser with complementary capabilities that we use less often. Using both browsers together was an interesting experience for us, sometimes uncomfortable, sometimes pleasant, but yielding useful insights.

On the uncomfortable side, we sometimes experienced frustration when we wanted to do a thing in our non-default browser that, to us, feels fluid and friction-free in IGB thanks to our deep, expert knowledge of how IGB works and what it can do. Indeed, our advanced expertise in our default browser seemed to magnify this discomfort.  An example was when we wanted to “zoom in” to inspect and count the base pairs separating two nearby donor sites supported in the data, a task that to us feels easy and fluid in IGB.

On the fun and exciting side, we learned more about the gene than we would have done otherwise. It was gratifying to use newer data to answer an old question that was previously impossible to address. It was also fun to explore the UCSC browser and observe how it organizes and describes information.  As we became familiar with its current user interface and display terminology (e.g., the “squished” track display mode), we developed a new appreciation for how it approaches the same problems we also have had to solve, such as how to communicate capabilities. We also appreciated the attention paid to ensuring users can look up the source of all data tracks, which is essential to understanding and interpreting what we see.
